# The one that abstained: *Psilocybe fuscofulva* genome suggests two recent origins of the psilocybin gene cluster in *Psilocybe*

**DOI:** 10.64898/2025.12.30.697041

**Authors:** Jason Slot, Alexander Bradshaw, Bryn Dentinger, Jan Borovička, Zachary Konkel, Alan Rockefeller, Ian Bollinger

## Abstract

Production of the psychoactive compound psilocybin is a defining feature of the genus *Psilocybe*, commonly referred to as “psychedelic mushrooms”. However, *Psilocybe fuscofulva* is a striking exception within *Psilocybe sensu stricto* as it lacks the stereotypical blue bruising characteristic of the genus, and psilocybin has not been detected in the species.To investigate the evolutionary events leading to differential psilocybin production among *Psilocybe* species, we produced genome assemblies two *P. fuscofulva* strains, one *Psilocybe polytrichoides* strain, and one *Psilocybe tampanensis* strain, complemented by reannotated public genomes and metagenome-derived assemblies from fungarium specimens. This sample represents both major *Psilocybe* clades (Clade I and Clade II) and the most closely related genera. Phylogenomic analysis based on 100 single-copy orthologs curated for high branch support strongly placed *P. fuscofulva* as the earliest-diverging lineage in *Psilocybe* Clade I. No psilocybin gene cluster (PGC) homologs, whether clustered or dispersed, were identified in *P. fuscofulva*, whereas a single intact PGC was present in all other examined *Psilocybe* genomes. The PGC resides in two distinct, clade-specific genomic loci: one conserved in Clade I and another in Clade II, each displaying characteristic gene orders and orientations consistent with rearrangement through circular intermediates. Time-calibrated phylogenies estimated the *Psilocybe* crown group at approximately 28 million years ago, with major clade divergences occurring in the Miocene. The absence of the PGC in *P. fuscofulva*, together with clade-specific structural conservation and the lack of remnant sequences at alternate loci, supports two independent origins of the PGC within *Psilocybe*: one in the ancestor of Clade II and a subsequent origin in Clade I following divergence from *P. fuscofulva*, most likely via horizontal gene transfer (HGT). Gene phylogenies provide weak support for transfer from Clade I to Clade II, although broader sampling is required for confirmation. These results constrain the timeframe of PGC emergence and dispersal to the Miocene, implying rapid HGT events possibly driven by ecological pressures in expanding grassland ecosystems. This study challenges the assumption of an ancestral psilocybin pathway in *Psilocybe* and its close relatives and underscores multiple recent acquisitions of the PGC that suggest it is an ecologically important metabolic trait in psychoactive fungi.

## Introduction

The mushroom-forming genus *Psilocybe* is known for its species’ ability to produce the psychedelic substance, psilocybin. Monographs by Guzmán (Guzmán 1983; 1995) provide the authoritative foundation for the taxonomy of *Psilocybe* based on morphological features, which have been refined recently by molecular phylogenetics (Moncalvo et al 2000; Borovicka et al 2015; Ramirez-Cruz et al 2013; Bradshaw et al 2024a). Before molecular phylogenetics, *Psilocybe* was conflated with psilocybin-absent species that have been transferred to other genera, especially with the historically “non-psychoactive” *Psilocybe,* now known to be more distantly related as the genus *Deconica* and *Leratiomyces* (Redhead et al 2007). Reclassification of these psilocybin-absent species is ongoing (Cao et al 2025; Bradshaw et al 2024a; Ramirez-Cruz et al 2020), with *Psilocybe* sensu stricto retaining the psilocybin-containing species (Redhead et al 2007). Recent phylogenomic analyses within *Psilocybe* s.s. have revealed strong support for two distinct clades (Bradshaw et al 2024a). All described members of Clade II, including the most widely cultivated species, *P. cubensis* (McTaggart et al 2023), produce psilocybin (Bradshaw et al 2024a). Similarly, Clade I species, which includes the type species, *Psilocybe semilanceata* (a.k.a. “Liberty Cap”), and *Psilocybe mexicana,* from which psilocybin was first isolated and structurally elucidated (Hoffmann et al 1958), also all produce psilocybin. However, *P. fuscofulva,* formerly referred to as *Psilocybe atrobrunnea* or *Psilocybe turficola,* is the only known species of *Psilocybe* s.s. that does not produce psilocybin. This species has remained an enigma as previous systematic studies using three DNA barcoding loci recovered it as an early branching species without strong placement in either Clade I or Clade II (Borovička et al 2015; Gotvaldová et al 2022). The lack of psilocybin in *P. fuscofulva* is an anomaly within *Psilocybe* s.s. and provides an opportunity to further explore the genetic basis and evolution of psilocybin biosynthesis.

Psilocybin production has been shown to be encoded by gene clusters with homologs in the saprotrophic genera *Psilocybe*, *Gymnopilus*, *Panaeolus*, *Pluteus*, and *Pholiotina*, and an independently evolved cluster is proposed to be responsible for psilocybin production in the ectomycorrhizal genus *Inocybe* (Fricke et al 2017, Reynolds et al 2018, Awan et al 2018, Bradshaw et al 2024; Schäfer et al 2025). The limited occurrences of psilocybin gene clusters (PGC) in the distant genera *Panaeolus*, *Pluteus*, and *Pholiotina* were inferred to have originated by horizontal gene transfer (HGT) from *Psilocybe* and *Gymnopilus* (Awan et al 2018; Reynolds et al 2018). While no clear mechanism for these HGT events has been proposed, alternate gene orders in each genus are consistent with a circular intermediate state (Awan et al 2018), which is characteristic of the mechanism of some mobile elements in fungi (Bucknell & McDonald 2023). The PGCs in *Psilocybe* and *Gymnopilus* by contrast might be assumed to have emerged in their common ancestor because of their phylogenetic proximity (Meyer & Slot 2023). However, because there are no intermediate forms of the PGC and unclustered homologs of its genes are distantly related, the timing and process of its origin remain unclear. Prior phylogenomic analyses inferred an ancestral origin of the psilocybin biosynthetic gene cluster (PGC) in *Psilocybe* ∼67 million years ago but lacked genomic data from closely related genera and the non-producing *P. fuscofulva*, which can skew molecular dating analysis (Bradshaw et al 2024a). While *P. fuscofulva* has been weakly placed as an early-diverging lineage in previous studies, it has never been investigated at the genome scale.

Resolving the phylogenetic placement and genetic composition of *P. fuscofulva* is key to reconstructing the origin and evolution of psilocybin biosynthesis in *Psilocybe* because of its *P. fuscofulva* lacks psilocybin production and holds an ambiguous position at the base of *Psilocybe* s.s. Therefore, we set out to generate new genomic sequence data to support the precise phylogenetic placement of *P. fuscofulva* in *Psilocybe* and to determine the existence and clustered state of the PGC in *P. fuscofulva.* To account for geographic variability, we sequenced the genomes of two strains, one fungarium specimen from the Czech Republic, *P. atrobrunnea* (= *P. fuscofulva,* PRM-922256) (Borovička et al 2015) and one pure culture from New York (Mushroom Observer #471082). We compiled additional genomes from fresh culture, fungaria vouchers, and public repositories to expand represention of the diversity of Clade I and Clade II, and the immediate relatives of *Psilocybe* for a more robust evolutionary assessment of the PGC.

Here, we resolve *P. fuscofulva*’s placement using high-quality genomes and infer that the PGC was independently acquired in each major *Psilocybe* clade via HGT. Through analysis of whole genome content, gene order, PGC location and structure, species phylogenies, we placed *P. fuscofulva* as the earliest branch of Clade I and inferred that it never had the PGC. Phylogenies of PGC genes weakly place Clade II within Clade I when rooted with *Gymnopilus*. Together, these results lead us to propose that there was one recent origin of the PGC by HGT in *Psilocybe* Clade I following the divergence of *P. fuscofulva* and a second origin in *Psilocybe* Clade II by HGT from Clade I. This model compresses the timeframe for the origin and dispersal of this metabolic feature.

## Results

Four genomes were produced for this study, *P. fuscofulva* strain JCSPSIFUS1B derived from the spores of Mushroom Observer #471082, collected from moss/litter in Putnam Valley, NY, *P. fuscofulva* derived from the voucher specimen PRM-922256 collected in the Czech Republic, *P. (Galeropsis) polytrichoides* strain JCSGALPOL1A, derived from spores of Mushroom Observer #471082, collected on soil under pine in Eastern Shasta-Trinity National Forest, CA, and *P. tampanensis* strain JCSTAMDOCA, derived from spores of Mushroom Observer #481512, collected from grass in a commercial area in Lexington, SC. The inclusion of *P. polytrichoides* was opportunistic and allowed us to evaluate its suspected inclusion in *Psilocybe* s.s. The 43 Mb *P. fuscofulva* genome assembly produced for this study (N50 47.3 kb, largest contig 428 kb) comprises 12,878 predicted genes with a median gene length of 1.3 kb, and the 40.6 Mb *P. polytrichoides* assembly (N50 8.6 kb, largest contig 202 kb) comprises 13,278 predicted genes with a median length of 1 kb. Genome assemblies for isolated strains obtained from public databases and reannotated with the same parameters as *P. fuscofulva* were used for comparative analyses (Table 1; Supplemental Data S1). Additional data from five fragmented assemblies obtained from fungarium specimen metagenomes, which have been shown to produce discontiguous, yet useful quality genome assemblies (Bradshaw et al 2024a; Varga et al 2025; Dirks et al 2025), was used to enhance phylogenomic reconstructions.

**Table 1.** Genomes used in this study.

| Taxa* | Strain/collection ID | Source<sup>†</sup> | PGC<br>(+/-) | accession | Length<br>(Mb) | N50<br>(Mb) |
| --- | --- | --- | --- | --- | --- | --- |
| *Galerina marginata* | CBS 339.88 | M | - | GCA_000697645.1 | 59.42 | 1.22 |
| *Ga. marginata* | ZP088764 | D | - | GCA_023014335.1 | 69.4 | 2.61 |
| *Ga. sulciceps* | ZP082096 | D | - | GCA_023014345.1 | 101.9 | 1.84 |
| *Gymnopilus dilepis* | SRW20 | M | + | GCA_002938385.1 | 45.5 | 0.035 |
| *Gy. junonius* | AH 44721 | N/A | - | GCA_015501075.1 | 59.5 | 0.152 |
| *Agrocybe pediades* | AH 40210 | N/A | - | GCA_015484485.1 | 45.1 | 0.105 |
| *A. praecox* | OKM6292 | N/A | - | GCA_964017145.1 | 46.4 | 0.350 |
| *Psilocybe cubensis* | FSU12409 | D | + | Psicub1_1 JGI | 40.5 | 0.136 |
| *P. cyanescens* | PBM2631 | M | + | GCA_002938375.1 | 47.5 | 0.055 |
| *P. serbica* | FSU12416 | D | + | Psiser1 JGI | 61.3 | 2.01 |
| *P. fuscofulva* * | JCSPSIFUS1B | D | - | SAMN54344997 | 49.9 | 0.041 |
| *P. fuscofulva* * | PRM-922256 | F | - | N/A | 59.9 | 0.008 |
| *P. mexicana* | FSU13617 | D | + | GCA_023853805.1 | 65.2 | 0.508 |
| *P. tampanensis* * | JCSTAMDOCA | D | + | PRJNA1395081 | 42.6 | 0.064 |
| *P. samuiensis* | WTU-F-055014 | F | + | N/A | 36.2 | 0.007 |
| *P. medullosa* | PRM-909630 | F | + | N/A | 81.7 | 0.006 |
| *P. weilii* | WTU-F-063525 | F | + | N/A | 67.4 | 0.011 |
| *P. stuntzii* | WTU-F-011520 | F | + | N/A | 37.5 | 0.068 |
| *P. zapotecorum* | iN: 36797577 | D | + | GCA_040207405.1 | 58.4 | 0.173 |
| *P. polytrichoides* * | JCSGALPOL1A | D | + | SAMN54344998 | 47.6 | 0.046 |

\*sequenced for this study
†M = Monokaryon culture; D = Dikaryon culture; F = Fungarium specimen

### Genomic data place *Psilocybe fuscofulva* at the base of Clade I

A phylogenomic tree (Figure S1) based on a dataset of 100 phylogenetically informative protein models (Supplemental Data S1) reproduced the previously reported topology of selected species. The key nodes supported [% bootstrap, gene concordance factor, site concordance factor] include monophyletic genera (*Galerina* [100, 100, 84.2], *Gymnopilus* [100, 99, 71.9], *Agrocybe* [100, 99, 64], *Psilocybe* [100, 99, 55.3]), *Psilocybe* Clade I (100, 88, 47.5) as previously defined, and *Psilocybe* Clade II (100, 97, 66.6). *P. fuscofulva* was placed sister to the previously defined Clade I [100, 80, 42.1]. Internal nodes in *Psilocybe* are consistent with previous phylogenies based on limited numbers of loci and genomes of fungarium specimens (Bradshaw et al 2024a; Bradshaw et al 2024b; Canan et al 2024; Van Der Merwe et al 2024; Ramírez-Cruz et al 2013; Ostuni et al 2024).

### The PGC is found in two alternate loci in *Psilocybe*

A single PGC was recovered from every *Psilocybe* genome except *P. fuscofulva*, where no evidence of any psilocybin genes was found (Figure 1). A single PGC was also recovered from *Gy. dilepis* as previously reported (Reynolds et al 2018), but only 3 PGC pseudogenes were recovered from *Gy. junonius* (Figure S2). There was no evidence of PGC genes in any other genomes in the dataset, namely *Galerina* and *Agrocybe* spp.

**Figure 1.**
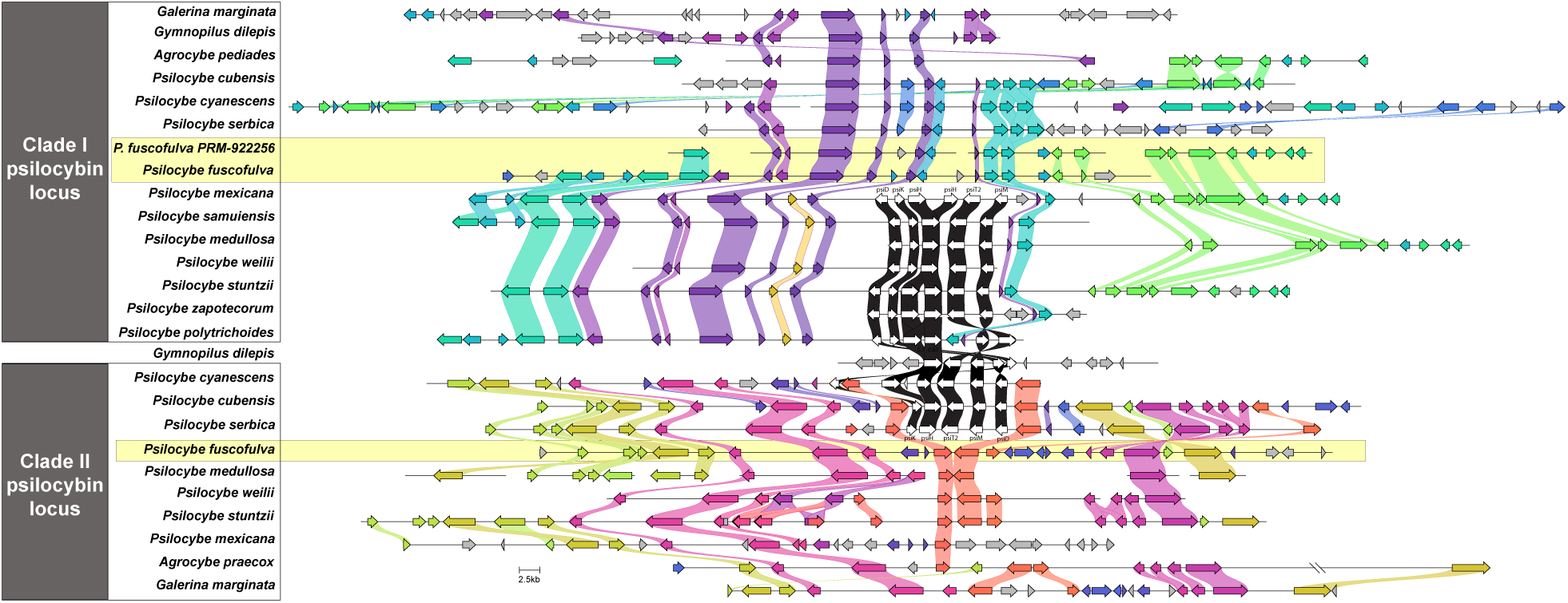
Two PGC loci in *Psilocybe*. The PGC (white arrows) is shown in two loci, one conserved in *Psilocybe* Clade I and the other conserved in *Psilocybe* Clade II. Conservation of synteny is indicated by common-colored gene sets and connecting lines. The PGC in *Gy. dilepis* is located in an unrelated locus. Homologous loci in *P. fuscofulva*, which lack PGC sequence, are highlighted in yellow. No PGC homologs or fragments thereof were detected in *P. fuscofulva* scaffolds after extensive searches. *P. zapotecorum* (Miller et al. 2024) was used in this figure only.

The order and orientation of PGC genes is generally specific to each *Psilocybe* clade, with some modifications present within each clade. The gene order and orientation in Clade I generally follows *psiD* (R), *psiK* (F), *psiH* (F), *psiT2* (R), *psiM* (R). Head to tail tandem duplications of *psiH* are present in *P. mexicana*, *P.* stuntzii, P*. zapotecorum*, and *P. polytrichoides*. The PGC in *P. polytrichoides* has a further alteration in gene order consistent with one inversion between *psiT2* and the second flanking gene of *psiM* that brought two flanking genes into the cluster between *psiH* and *psiM*, and a second inversion of *psiM* only. The gene order and orientation in Clade II generally follows *psiK* (F), *psiH* (F), *psiT2* (R), *psiM* (R), *psiD* (R). A head to tail tandem duplication of *psiH* is present in *P. cyanescens*. The PGC in *P. cyanescens* has a further alteration in gene order consistent with an inversion from one flanking gene of *psiK* to a third tandem duplicate of *psiH* that brought a gene provisionally named *psiT1* by Fricke et al (2017) into the cluster. The alternate primary gene orders in Clade I and Clade II are consistent with a circular intermediate state between the two. The PGC gene order and orientation in *Gy. dilepis* is *psiH* (F), *psiT2* (R), *psiM* (R), *psiD* (F), *psiK* (F). This is consistent with an inversion of *psiD* and a circular intermediate between each *Psilocybe* clade PGC. These clade-specific rearrangements are consistent with independent cluster insertions via circular intermediates following separate HGT events.

PGCs in *Psilocybe* were found in one of two loci. In Clade I species, the PGC occupies one conserved locus. The psiD flank features a putative hypothetical protein, transaldolase, and p-nitrophenyl phosphatase. The psiM flank includes a putative 60s ribosomal protein, hypothetical protein, and capsular-associated protein. In Clade II species, the PGC is located in a second locus characterized by a putative *alcE* dehydrogenase, *sc15* protein, and PPR repeat family protein on the *psiD* flank and a putative helix loop helix protein, two hypothetical proteins, and a major facilitator superfamily transporter previously annotated as *psiT1*. Searches for remnant sequences of PGCs in the Clade I locus of Clade II species and in the Clade II locus of Clade I species recovered no evidence of prior presence at the respective alternate loci. *P. fuscofulva* and outgroup taxa in *Gymnopilus*, *Agrocybe*, and *Galerina* lacked evidence of any PGC at either the Clade I and the Clade II loci. Alignments of genomic scaffolds across all taxa demonstrate conservation of gene order in sequence adjacent to the PGC in the Clade I and Clade II loci, with the exception of the PGC, which is only present in the respective loci in Clade I and Clade II *Psilocybe*. Both loci in *P. fuscofulva* resemble homologous loci in related genera. The PGC is at an alternate locus in *Gy. dilepis* and pseudogenes of *psiD*, *psiK*, and *psiM* are each on a different, unrelated scaffold in *Gy. junonius*.

### Molecular clock analysis suggests an Oligocene origin of *Psilocybe* and younger PGC

A time tree (Figure 2) based on a secondary calibration of the basal divergence time derived from previous studies (Ruiz-Dueñas et al 2021; Varga et al 2019) placed the origin of *Psilocybe* at 27.65 Ma (CI: 17.495-43.6997 Ma). *P. mexicana* was substituted with the very closely related *P. tampanensis* (Bradshaw et al 2024a) because, while the assembly of the PGC was incomplete due to a lack of longer sequence reads, individual gene sequences were more reliable due to depth of Illumina sequence reads.

**Figure 2.**
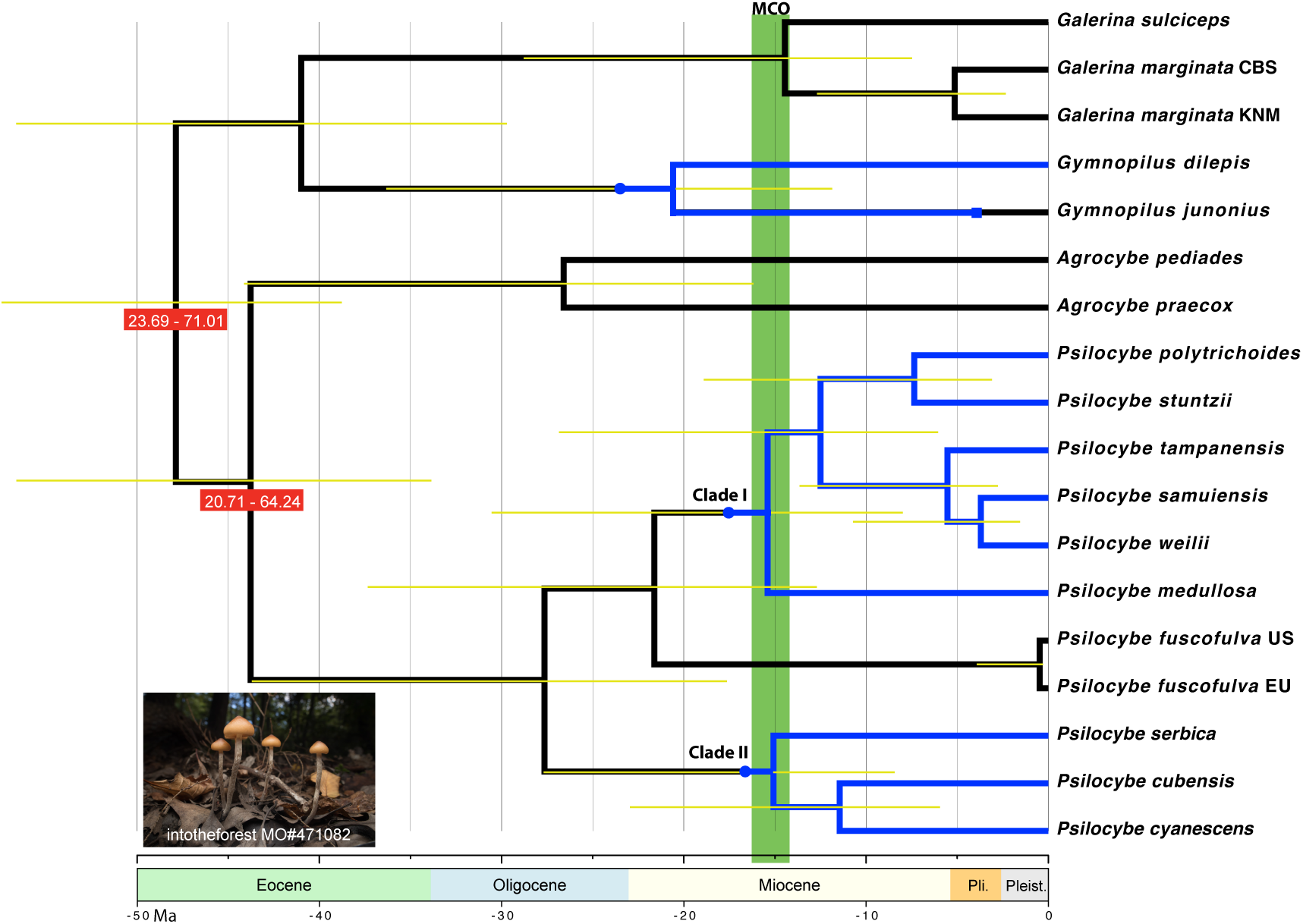
Time tree of genomes used in this study based on two secondary calibration ranges. A hypothetical reconstruction of PGC presence is indicated by blue branches with round points indicating putative origins and a square point indicating loss of function. Calibration ranges are indicated with red boxes and posterior confidence intervals with yellow bars. Photo inset courtesy of intotheforest (CC BY-SA 3.0) is the collection (Mushroom Observer #471082), from which *Psilocybe fuscofulva* JCSPSIFUS1B was isolated.

Time-calibrated phylogenies were inferred under two calibration schemes: a single-calibration model (Supplemental Data S1) and a two-calibration model (Figure 2; Supplemental Data S1). Between the two schemes, age estimates from the two analyses were broadly congruent. Root ages differed by less than 1 Ma (47.2, single calibration, vs. 47.9 Ma, two calibrations). Under the single-calibration scheme, the most recent common ancestor (MRCA) of the ingroup was dated to 47.22 Ma (95% CI: 38.21–58.36 Ma), placing the origin of the lineage firmly in the Eocene. A subsequent deep split separating *Agrocybe* from *Psilocybe* was estimated at 43.17 Ma (95% CI: 33.34–55.91 Ma), also during the Eocene. Shortly thereafter, the divergence between *Galerina* and *Gymnopilus* from *Psilocybe* was inferred at 40.43 Ma (95% CI: 29.23–55.92 Ma), suggesting that the major generic lineages were established prior to the end of the Eocene.

Within *Psilocybe*, the primary split between Clade I and Clade II occurred during the late Oligocene, dated to 27.27 Ma (95% CI: 17.26–43.10 Ma). Diversification within *Agrocybe* was inferred at a similar time, with the split between *A. pediades* and *A. praecox* dated to 26.23 Ma (95% CI: 15.81–43.53 Ma). Early Miocene diversification within *Psilocybe* included the separation of *P. fuscofulva* from the psilocybin-producing Clade I at 21.32 Ma (95% CI: 12.36– 36.79 Ma), and the divergence between *Gy. dilepis* and *Gy. junonius* at 20.30 Ma (95% CI: 11.52–35.78 Ma). Subsequent Miocene diversification involved multiple internal splits within both *Psilocybe* clades. Within Clade I, a split separating *P. medullosa* from other Clade I taxa was dated to 15.19 Ma (95% CI: 7.68–30.05 Ma), followed by further internal divergences at 12.33 Ma (95% CI: 5.76–26.39 Ma), 7.25 Ma (95% CI: 2.83–18.55 Ma), and 5.46 Ma (95% CI: 2.28–13.06 Ma). Within Clade II, the divergence between coprophilic (*P. cubensis*) and wood-decaying (*P. cyanescens*) lineages was dated to 11.29 Ma (95% CI: 5.66–22.53 Ma). Diversification within *Galerina* continued into the Miocene, with the split between *Ga. Sulciceps* and two *Ga. marginata* genomes dated to 14.25 Ma (95% CI: 7.18–28.31 Ma).

Shallow divergences extended into the Pliocene and Pleistocene. The split between the two *Ga. marginata* genomes was inferred at 5.07 Ma (95% CI: 2.08–12.35 Ma), while diversification within Clade I continued into the Pliocene with the separation of *P. samuiensis* and *P. weilii* at 3.65 Ma (95% CI: 1.31–10.17 Ma). The most recent divergence in the dataset involved the North American and European *P. fuscofulva* genomes, dated to 0.46 Ma (95% CI: 0.06–3.54 Ma), consistent with a very recent Pleistocene diversification.

Incorporating a second calibration point yielded a highly similar temporal pattern. Root ages differed by less than 1 Ma (47.88 Ma, 95% CI: 38.74–59.17 Ma), and most internal nodes shifted by approximately 0.3–0.6 Ma, without altering the relative ordering of divergence events. Across both calibration schemes, results consistently support an Eocene origin of the ingroup, late Oligocene establishment of major *Psilocybe* clades, and Miocene-to-Pliocene diversification within clades, indicating that the inferred evolutionary timeline is robust to calibration strategy (Supplemental Data S1).

### PGC gene phylogenies place Clade II species in Clade I

Phylogenetic analyses of PGC amino acid sequences (Figure S3) are consistent with similarity of gene order within clades. However, because there are only three clades of PGCs represented and no close outgroups PGC genes are known, gene trees provide limited information about the origin and dispersal within each clade. Homologous sequences from *Athelia termitophila*, previously suggested as the closest relatives of those in the PGC (Reynolds et al 2018) were not considered an acceptable root due to excessive divergence, varied copy number, and dispersal among multiple gene clusters (Konkel et al 2021). Four sequences (PsiD, PsiM, PsiT2, and PsiK) weakly place the Clade II *Psilocybe* PGC within Clade I *Psilocybe* when rooted with *Gy. dilepis*. In PsiD (62%) and PsiM (51%), Clade II is weakly supported as sister to *P. medullosa*, in PsiT2 (69%) and PsiK (68%), Clade II is weakly supported as sister to the remaining Clade I taxa.

Clade I is monophyletic with no support (42%) in PsiH. Phylogenetic analysis also suggests PsiH duplicates within Clade I and Clade II are respectively monophyletic, consistent with convergent duplication events following the independent origin of the cluster in each clade. The duplication in Clade I occurred after the divergence of *P. medullosa* and was not retained in the common ancestor *P. samuiensis* and *P. weilii.* Independent duplications in Clade II resulted in multiple paralogs in both *P. serbica* and *P. cyanescens*.

## Discussion

*Psilocybe fuscofulva* is the only species of *Psilocybe* s.s. for which studies have not detected psilocybin/psilocin (Borovička et al 2015). Our phylogenomic analysis strongly suggests this species is deeply divergent and the earliest branch of Clade I. The absence of any canonical psilocybin biosynthesis genes either dispersed or clustered within the *P. fuscofulva* genome contributes strong evidence that this species also lacks the genetic capability of synthesizing psilocybin. Together with further evidence that the PGC has two conserved gene structures that correspond to the reciprocally monophyletic Clades I and II, we propose that there were two asynchronous, independent origins of the PGC within *Psilocybe*. The phylogenies of PGC genes overall suggest HGT from Clade I to Clade II, but support is weak and additional potential donor taxa cannot be excluded until *Psilocybe* PGCs can be shown monophyletic (except for the previously reported horizontal transfer to *Panaeolus* [Awan et al 2018; Reynolds et al 2018]) with further sampling in *Gymnopilus* and other genera.

While it has regularly been assumed that the production of psilocybin was to some extent an ancestral trait, in the ancestor of *Gymnopilus* and *Psilocybe* or at least a synapomorphy of the genus *Psilocybe*, our results imply there were instead multiple recent origins. If the PGC is not ancestral to *Psilocybe*, it cannot be inferred to be ancestral to the common ancestor of *Gymnopilus* and *Psilocybe*. Instead, we reconstruct an origin of this PGC between ∼20 and ∼25 Ma with multiple HGT events occurring in the ensuing 5-10 million years. The current study’s Miocene (∼27-28 Ma) dual origins contrast sharply with the Bradshaw et al (2024a) estimate of a single ∼67 Ma origin. This compression of the timeline of origin and dispersal of the PGC raises questions about what ecological factors may have been driving and facilitating its dispersal.

Acknowledging the uncertainty inherent to molecular dating analyses, this range roughly overlaps with the Miocene Climate Optimum (MCO), a period of time inferred to be much warmer globally than today, along with the major expansions of grasslands where novel niches and pressures emerged for mushroom-forming fungi (Edwards et al 2010; Strömberg 2011).

Such niches include the decay of grass, herbivore dung, and other soil biomass in grasslands. Yet psilocybin production may relate more to pressures from mycophagists like grassland rodents, insects, and mollusks that simultaneously adapted to and diversified in these environs (Bradshaw et al 2024; Elliot et al 2022; Kitabayashi et al 2022; Santamaria et al 2023).

Reconstructing the gain and loss of traits millions of years ago is difficult to achieve unambiguously. It is possible, but less parsimonious, that *Psilocybe* s.s. had a single gain of the PGC ∼28 Ma, followed by a loss in an ancestor of *P. fuscofulva*, with or without a regain of the PGC in Clade I. In this scenario, major rearrangement and relocation would need to have occurred in the MRCA of either Clade I or Clade II species (or both). Given the high conservation of gene order within each of the respective clade PGCs that has persisted for millions of years, scenarios that require invoking evolutionarily vertical cluster rearrangement seem less likely. Genomic data exist only for a relatively small number of psilocybin-producing mushrooms, biased to *Psilocybe* s.s., and genomic data from their closest relatives are minimal. Distinguishing between these two scenarios requires broader genomic sampling and evidence of the mechanism underpinning HGT of the PGC in mushrooms.

Although our phylogenetic placement of *P. fuscofulva* received strong statistical support and resolves inconsistent earlier results from a three gene analysis (Borovička et al 2015), the possibility remains that its placement is erroneous. Importantly, the placement of *P. fuscofulva* in or sister to Clade I received a gene concordance factor of 80% but with an average site concordance factor of 42.1%, representing less than half of all sites decisively supporting its position. The site concordance factor for Clade I and the genus *Psilocybe* are similarly low at 47.5%, and 55.3% respectively. This suggests there may be underlying gene conflict that needs to be explored with more extensively sampled data sets, both regarding genes and species diversity.

While the directionality and mode of HGT cannot be reliably determined from these data, these results bring further attention to previous observations that PGC HGT events are associated with rearrangements consistent with circular intermediates (Awan et al 2018). Plasmid-like intermediates are known in HGT in other eukaryotes, such as in the transfer of mitochondrial DNA tracts in the holoparasitic plant *Lophophytum mirabile* (Roulet et al 2024) and the transfer of nuclear DNA fragments in the form of extrachromosomal circular DNA (eccDNA) in grafts of tomato plants on goji stock (Zhang et al 2024). Moreover, growing evidence highlights a significant role for large mobile genetic elements (a.k.a. "Starships"), including a proposed circular intermediate, as demonstrable agents of horizontal gene transfer (HGT) in Ascomycota (Bucknell & McDonald 2023; Urquhart et al 2024). This raises the possibility PGCs may have been transferred via an ancient "starship" with limited evidence remaining (but see tyrosine recombinase in PGCs, Bollinger et al 2025). However, transposable elements analogous to starships have not yet been characterized in Basidiomycota. We present a reconstruction of hypothetical HGT events involving circular intermediates (Figure S4) while remaining agnostic with regards to the origin of the PGC, which would enable arranging this model in a phylogeny, because the currently inadequate sample of genomes, lack of intermediates in PGC assembly, and uncertainty in gene trees leave multiple hypotheses possible. Broader sampling of *Gymnopilus* and outgroups is needed to root PGC phylogenies robustly and confirm transfer directionality. To our knowledge, the PGC represents one of only three reported horizontally transferred gene clusters in Agaricales (Konkel et al 2024; Luo et al 2022; Reynolds et al 2018), suggesting that strong selection may have been a key factor in its HGT. This is reinforced by the high conservation of the PGC in a background of low rates of gene clustering in Agaricomycetes (Konkel 2023). Physical transfer of these genetic elements may be mediated through hyphal contact, or by third parties such as mycophagous animal vectors, bacteria, or viruses.

## Methods

### Strain cultivation

Spores from *P. fuscofulva* (Mushroom Observer #471082), *P. polytrichoides* (Mushroom Observer #367492), and *P. tampanensis* (Mushroom Observer #481512) were germinated on 1/4 strength potato dextrose broth, and fast-growing colonies were isolated and presumed to be dikaryotic.

### Fungarium samples

Specimens from fungaria were chosen to expand sections of both Clade I and Clade II. Specimen metadata was accessed through either MycoPortal (Miller & Bates 2017) or directly through a respective holding institutions online collection database. Full Specimen loans or sampled gifts were requested for each specimen in compliance with the Nagoya protocol and Convention on International Trade in Endangered Species of Wild Fauna and Flora (CITES) when applicable. Index Herbariorum (Thiers 2023) codes and collection catalogue numbers are reported in Table 1.

### DNA extractions and sequencing

*P. fuscofulva* JCSPSIFUS1B was cultured on liquid rich media consisting of trace salts from concentrated stock solutions (CaCl₂·2H₂O at ∼0.5 mL/L, FeCl₂·6H₂O at ∼0.5 mL/L, NaCl at 1 mL/L, MgSO₄·7H₂O at 1 mL/L, (NH₄)₂HPO₄ at 5 mL/L, and KH₂PO₄ at 10 mL/L), multiple carbon sources (2 g/L malt extract, 5 g/L potato dextrose broth, 5 g/L dextrose, and 2 g/L cellobiose), nitrogen sources (2 g/L polypeptone peptone and 1 g/L yeast extract), post-autoclaving addition of 1× filter-sterilized BME vitamin solution (from a 100× frozen stock aliquot). *P. tampanensis* JCSTAMDOCA and *P. polytrichoides* JCSGALPOL1A were cultured in potato dextrose broth at room temperature for 30 days without shaking. Tissue was washed through Miracloth with sterile MilliQ water and pulverized with a mortar and pestle in liquid nitrogen. Genomic DNA was immediately extracted using the DNeasy plant minikit (Qiagen). Short-read DNA libraries were prepared using the NEBNext Ultra DNA library preparation kit and were sequenced with a 150-bp paired-end format on a NovaSeq 6000 system (Illumina) by Novogene.

gDNA samples of fungarium specimens were prepared following the phenol-chloroform extraction methodology as Bradshaw et al 2024. In brief ∼5-15 mg of dried Hymenophore fragments were mechanically homogenized in 2.0 mL screw-cap tubes containing a single 3.0- mm and 8 × 1.5-mm stainless steel beads and shaking them in a BeadBug™ microtube homogenizer (Sigma-Aldrich, #Z763713) for 120 s at a speed setting of 3,500 rpm. Following physical homogenization, lysis was performed using 500 ml of tissue lysis buffer from the Monarch Genomic DNA Purification Kit (NEB, #T3010S) with 10ul Proteinase K (∼20mg/ml) incubated at 56C and 800RPM on a thermal mixer overnight. After chemical lysis, 7ul of RNase A (∼10mg/ml) was added and incubated at 56C and 800RPM for one hour. gDNA was then purified using a phenol:chloroform:isoamyl (24:25:1) protocol with homemade phase lock tubes containing high vacuum grease (Molykote, ASIN:B0DGCFHP9B), a 1:1 mixture of phenol:chloroform:isoamyl and sample were rotated on a hula mixer for ten minutes, then spun down at maximum speed (>12,000G) in a microcentrifuge. This step was repeated twice. gDNA was then precipitated using 5M NACL to a final concentration of 0.3M with double the amount of absolute ethanol, washed in cold 70% ethanol twice, and eluted in 150ul of Monarch DNA Elution Buffer (T1016-3). Finally, 75ul of the gDNA samples were cleaned and concentrated using the Zymo Research DNA Clean & Concentrator -5 kit (D4013) with 5x the DNA binding buffer to account for the short fragment stereotypical of old fungarium vouchers. gDNAs were submitted to the High Throughput Genomics Core at the University of Utah, where sequencing libraries were prepared using the Nextera™ DNA Flex Library Prep (Illumina, #20018704) and sequenced on a full lane of Illumina NovaSeq 6000 PE 2 ×150 bp using an S4 flow cell.

### Genome assembly

Illumina reads of cultured *P. fuscofulva*, *P. tampanensis*, and *P. polytrichoides* were trimmed with Trimmomatic v0.36 (Bolger et al 2014) and assembled using SPAdes v3.12.0 (Bankevich et al 2012). *P. tampanensis*, and *P. polytrichoides* assemblies were later improved using the Entheome Genome Assembly Pipeline (EGAP) version 2.5 (iPsychonaut n.d.), executed utilizing sixteen (16) cpu-threads and sixty-four gigabytes (64GB) of RAM on a local Linux Subsystem within a core Windows operating system. EGAP integrates Oxford Nanopore Technologies (ONT) long-read and Illumina short-read sequencing data to assemble fungal and other eukaryotic genomes, leveraging multiple assembly algorithms (MaSuRCA version 4.1.2 [Zimin et al 2013], Flye version 2.9.5 [Kolmogorov et al 2019], & SPAdes version 4.0.0) while evaluating assemblies based on BUSCO odb12 databases for agaricales and basidiomycota (Simão et al 2015) using Compleasm version 0.2.7 (Huang & Li 2023) completeness and QUAST version 5.3.0 (Gurevich et al 2013) contiguity metrics (contig count, N50, L50, GC content). The pipeline automatically selects the optimal assembly using a composite score prioritizing BUSCO completeness (single + duplicated orthologs).

For fungarium specimen metagenomes, raw sequencing read quality control was performed using FastQC v 0.12.1 (Andersson et al 2010) and MultiQC v1.14 (Ewels et al 2016), then quality filtered and trimmed using FASTP version 0.23.4 (Chen et al 2018). Genome assembly was performed using metaSPAdes v3.15 (Nurk et al 2017) with a custom kmer profile of 21, 77, 85, 99, 111, and 127, as it was found that this profile favored overall better contiguity using larger kmers with the added high accuracy of small kmer values. Post processing of the assemblies was performed using Redundans v 2.01 (Pryszcz & Gabaldón 2016) to reduce heterozygosity and improve contiguity for better gene prediction downstream. Genome completeness was measured using Benchmarking Universal Single-Copy Orthologs (BUSCO) v 6.0.0 (Simão et al 2015) with the AUGUSTUS v3.14 (Stanke et al 2006a) gene prediction pipeline using the coprinus_cinereus species training set and the agaricales_odb12 and basidiomycota_odb12 reference database. Assembly statistics were performed using QUAST v 5.3.0 (Gurevich et al 2013) (Supplemental Data S1).

#### Genome annotation

Genome assemblies of *P. fuscofulva, P. tampanensis, P. polytrichoides*, metagenome-derived assemblies, and select strains obtained from public databases were annotated by a common pipeline based on Funannotate 1.8.17 (Palmer and Stajich 2020). The assemblies were first cleaned by removing contigs shorter than 500bp and soft-masked using the ‘mask’ command. A preliminary BUSCO database was created with the funannotate-BUSCO2.py script using the basidiomycota_odb9 lineage dataset. Gene prediction was then performed with the ‘predict’ command in Funannotate, which used Augustus, GlimmerHMM, SNAP, and GeneMark-ES (Korf 2004; Majoros et al 2004; Stanke et al 2006b; Borodovsky & Lomsadze 2011). *P. cubensis* (GCF_017499595.1) transcript evidence was aligned to the assembly with minimap2 (Li 2018) and published annotations of *P. serbica, P. cyanescens, P. cubensis, Panaeolus papillionaceus, Panaeolus cyanescens, Gy. junonius, Gy. dilepis, Agrocybe pediades, Cyclocybe aegerita* and UniProt/Swiss-Prot were used as protein evidence for AUGUSTUS predictions. EvidenceModeler was used to filter best predictions by giving equal weight to prediction methods and double weight to high-quality predictions extracted by AUGUSTUS (Hoff & Stanke 2019).

### Phylogenomic analysis

To standardize gene predictions, all annotations except those derived from metagenomes were amended using a fork of orthofiller (Dunne & Kelly 2017; xonq n.d.). Orthofinder v2.2.3 (Emms & Kelly 2019) was used to identify single copy orthogroups (SCO). SCOs were randomly sampled and analyzed until 100 trees with an average bootstrap percentage of at least 80 were identified. Protein sequences were aligned with MAFFT v7.511(Katoh & Standley 2013), the alignment was trimmed with TrimAl v 1.4 (Capella-Gutiérrez et al 2009), and the gene tree inferred using RAxML-NG v. 1.2.0 (Kozlov et al 2019). Nearest orthologs of each SCO were then retrieved from metagenome-derived assemblies using Exonerate v.2.2.0 v.2.2.0 (Slater & Birney, 2005) with the protein2genome:bestfit match of the *P. fuscofulva* ortholog. The phylogenomic tree was inferred using a concatenated matrix of all 100 SCOs in IQ-TREE v.2.2.2 COVID-edition (Minh et al 2020) with 1000 ultrafast bootstraps, gene concordance factors, and site concordance factors for support.

### Divergence time and molecular dating

Divergence times were estimated using the relative rate framework (RRF) implemented in RelTime within MEGA v11 (Tamura et al 2012; 2018; 2021). This approach was selected because it efficiently analyzes large phylogenomic datasets while providing divergence time estimates comparable in accuracy to Bayesian relaxed-clock methods such as BEAST (Drummond & Rambaut 2007) and MCMCTree (Yang & Rannala 2006) but with substantially reduced computational resources. Multiple sequence alignments were generated with MAFFT for the 100 single-copy loci previously identified for phylogenomic analysis. Molecular dating utilized the previously generated ML tree using *Psilocybe* s.s. as the ingroup and six selected outgroup taxa as the user-provided species tree for analysis (Figure S1). Amino acid substitution modeling followed the Whelan and Goldman + Freq. (WAG+F) model (Whelan & Goldman 2001). and branch lengths were estimated using maximum likelihood. Because deep fungal divergence dating can be sensitive to calibration placement (Berbee & Taylor 2010; Taylor & Berbee 2006), we analyzed the dataset under two calibration schemes. The first employed a single uniform prior applied to the node uniting *Psilocybe* and *Galerina* (MRCA of *P. cubensis* and *Ga. sulciceps*), with bounds set from published estimates (Min= 23.69 Max=71.01Ma) (Ruiz-Dueñas et al 2021; Varga et al 2019). In the second scheme, we applied uniform calibrations to two nodes: the MRCA of *Galerinaceae* and *Agrocybaceae* (*P. polytrichoides*–*A. praecox*) (Min=20.71, Max=64.24) and the MRCA of *Psilocybe* and *Galerina* (*P. cubensis*–*Ga. sulciceps*) (Min= 23.69 Max=71.01Ma).

### PGC evolutionary analysis

Genomic scaffolds containing homologs of were retrieved from each genome assembly using tblastn 2.10.0+ (Camacho et al 2009) and refined gene models were inferred using protein-guided alignment against the novel genome assembly using Exonerate with the "protein2genome:bestfit" model allowing 20% identity and maximum intron length of 300 bp using Tryptophan Decarboxylase (*PsiD*, UniProt: P0DPA6), Psilocybin-related N-methyltransferase (*PsiM*, UniProt: P0DPA9), Psilocybin-related Hydroxylase (*PsiH*, UniProt: P0DPA7), Psilocybin-related Phosphotransferase (*PsiK*, UniProt: P0DPA8), and Psilocybin-related Transporter (*PsiT2*, UniProt: P0DPB2) as queries. Flanking genes in scaffolds with manually confirmed PGC homologs within 10Kb of the PGC in Clade I and Clade II loci were predicted by blastx against GenBank followed by Exonerate using top hits as queries as previously described. Loci were aligned using clinker v.0.0.28 (Gilchrist & Chooi 2021) with exonerate gene coordinates and a minimum amino acid identity of 25%.

In order to analyze the phylogenies of PGC genes, amino acid sequences predicted with exonerate were aligned with MAFFT, the alignment was trimmed with TrimAl, and tree topologies were inferred using IQ-TREE v. 2.2.2 COVID-edition (Minh et al 2020) with 1000 ultrafast bootstraps, gene concordance factors, and site concordance factors for support with automatic model selection and 1000 ultrafast bootstrap replicates.

### Author Contributions

Conceptualized study – JCS, JB

Provided materials – AR, JB

Generated data – ZK, AB

Analyzed data – JCS, AB, IB

Writing – JCS, AB, BD, JB, IB, ZK

Figures – JCS, AB

### Data availability

All raw genomic sequencing data has been deposited in the Short Read Archive (SRA) under Bioproject numbers PRJNA1395081 and PRJNA1395118; Genome assembly and Biosample accession numbers are reported in Supplemental Data S1. All Type specimens derived molecular data have also been provided for RefSeq designation and curation.

Any code or specific script requests should be sent to the corresponding author.

## Acknowledgements

We would like to thank intotheforest, the Mushroom Observer contributor who collected the *P. fuscofulva* specimen, and Carlton Dunlap (Carlyd95), the iNaturalist contributor who collected the *P. tampanensis* specimen used for culturing. We would also like to thank the holding institutions who supported this work and provided material, much of which is rare and irreplaceable: The Natural History Museum of Utah (UT), University of Washington (WTU), and the National Museum of Prague (PRM). The work of JB was carried out as a part of the project MycoLife – World of Fungi under the Strategy AV21 program of the Czech Academy of Sciences.

---

**Figure S1.**
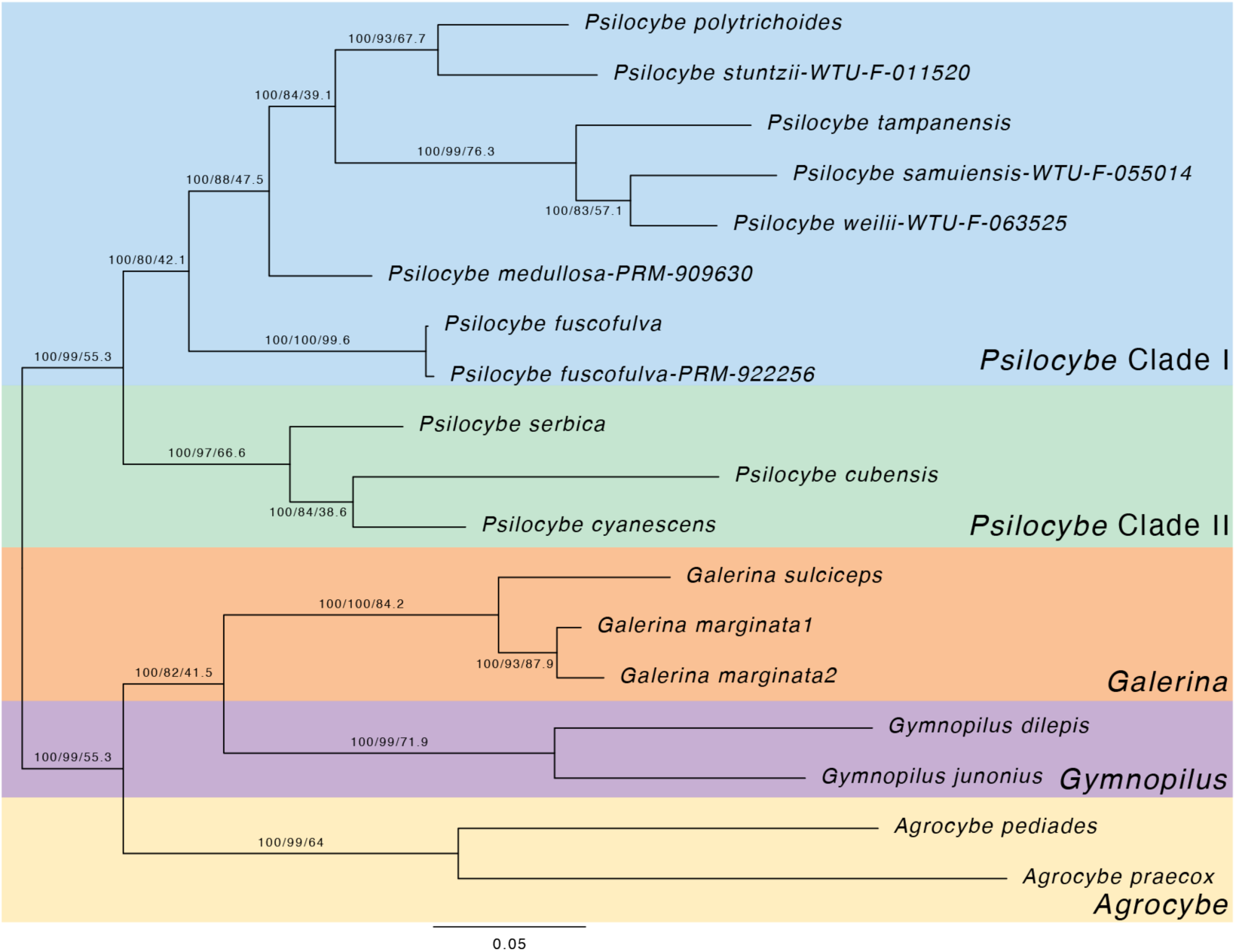
Phylogenomic analysis of the focal clade based on 100 single copy orthologs with a minimum of 80% average bootstrap support. Dataset was originally built with high quality pure culture assemblies and then supplemented by orthologs retrieved from filtered metagenomes of five fungarium specimens using exonerate. Phylogenomic analysis was performed using IQ-TREE 2.2.2 COVID-edition. Support values represent the percentage of 1000 ultrafast bootstraps in a concatenated gene analysis (left), gene concordance factor (middle), and site concordance factors (right) across all gene trees.

**Figure S2.**
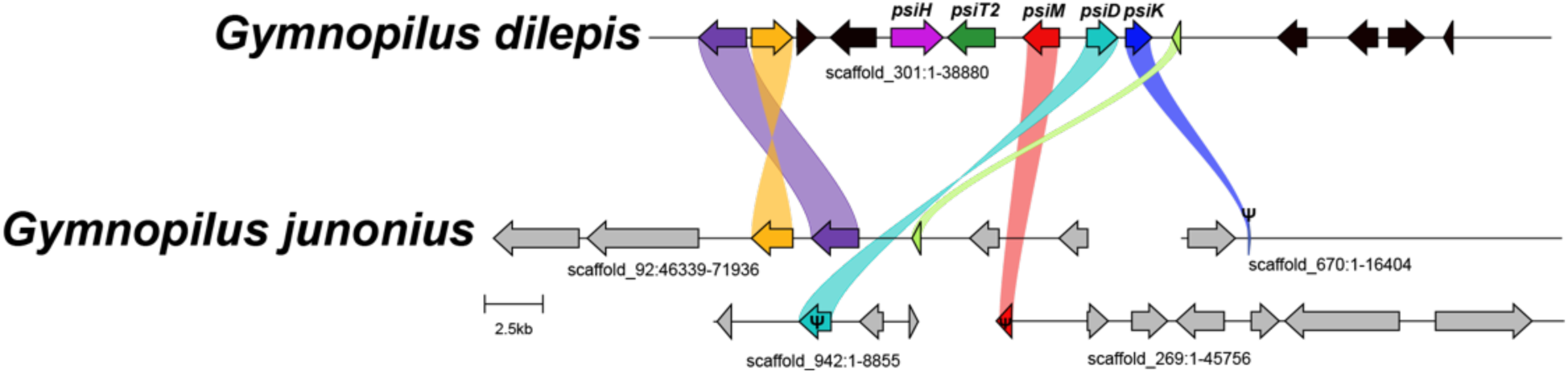
Evidence of loss of the PGC in *Gymnopilus junonius*. Three pseudogenes were recovered by tblastn and respective scaffolds were aligned using clinker and clustermap.

**Figure S3.**
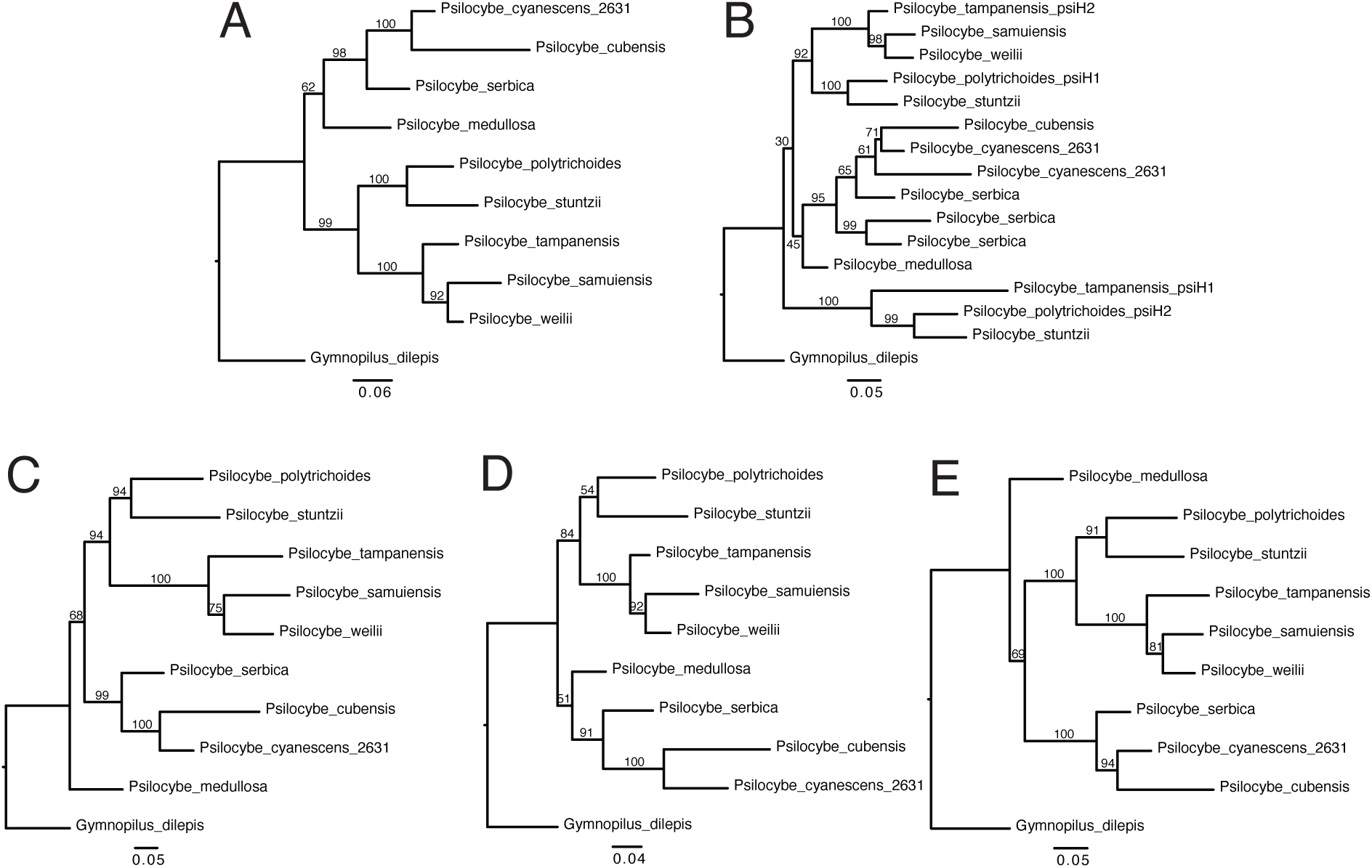
Phylogenetic analyses of PGC sequences weakly place Clade II within Clade. **I.** Maximum likelihood phylogenies of PGC amino acid sequences. A. PsiD, B. PsiH, C. PsiK, D. PsiM, E. PsiT2, with support values representing percentage of 1000 ultrafast bootstraps. All gene topologies are consistent with HGT from Clade I to Clade II of *Psilocybe*, but this assumes *Gy. dilepis* represents a true outgroup.

**Figure S4.**
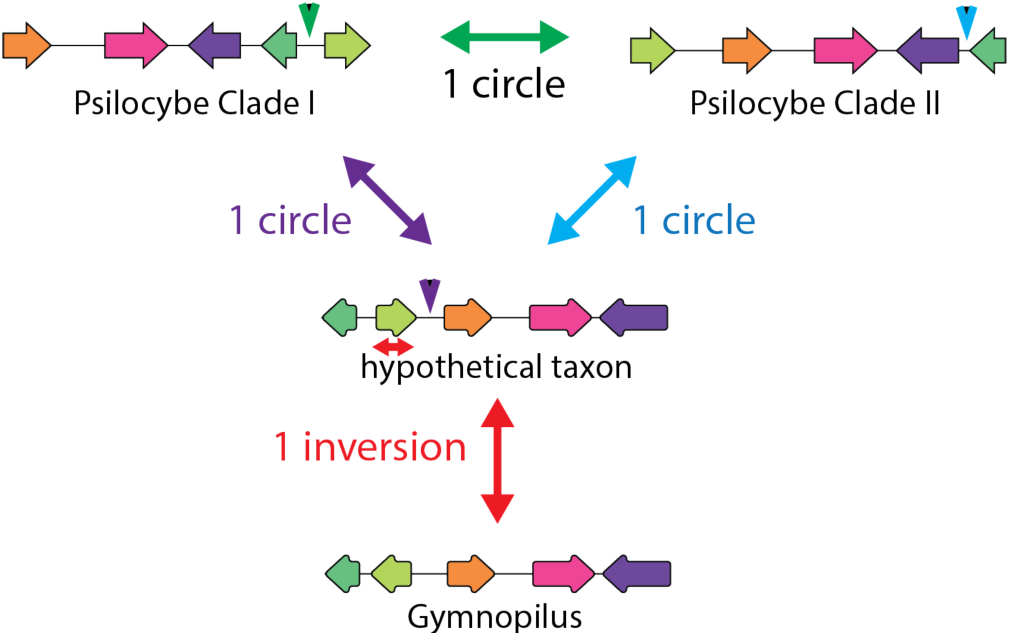
Molecular evolution events explaining the diversity of PGC gene order. One circular intermediate separates the gene order of *Psilocybe* Clade I, Clade II, and a hypothetical intermediate taxon, which differs from the PGC order in *<u>Gymnopilus</u>* by a single gene inversion. Breakpoints are indicated by arrowheads whose color corresponds to the circular intermediate, and the inversion is indicated by a bidirectional arrow.

