## Supplemental Figures for "The one that abstained: *Psilocybe fuscofulva* genome suggests two recent origins of the psilocybin gene cluster in *Psilocybe*"

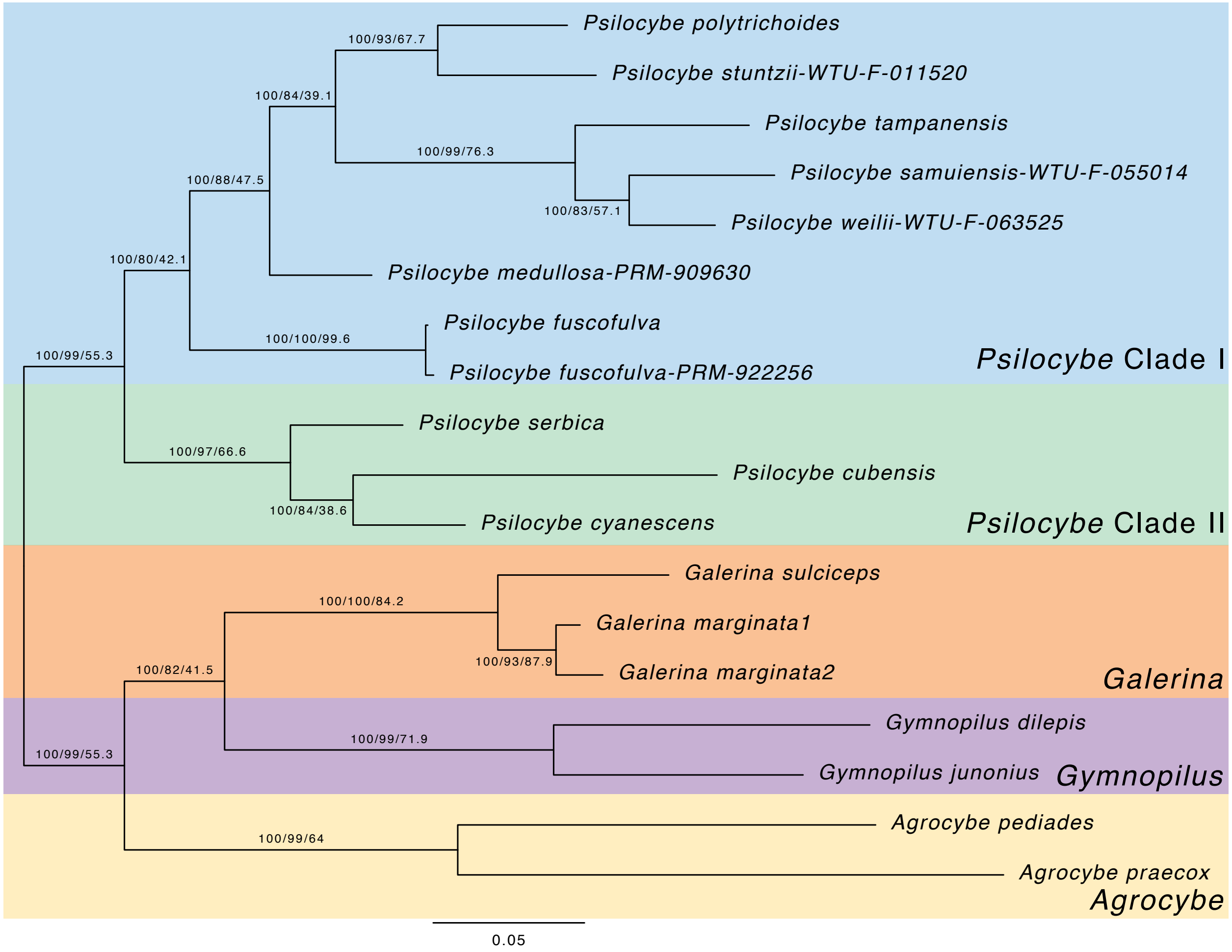


**Figure S1. Phylogenomic analysis of the focal clade based on 100 single copy orthologs with a minimum of 80% average bootstrap support.** Dataset was originally built with high quality pure culture assemblies and then supplemented by orthologs retrieved from filtered metagenomes of five fungarium specimens using exonerate. Phylogenomic analysis was performed using IQ-TREE 2.2.2 COVID-edition. Support values represent the percentage of 1000 ultrafast bootstraps in a concatenated gene analysis (left), gene concordance factor (middle), and site concordance factors (right) across all gene trees.

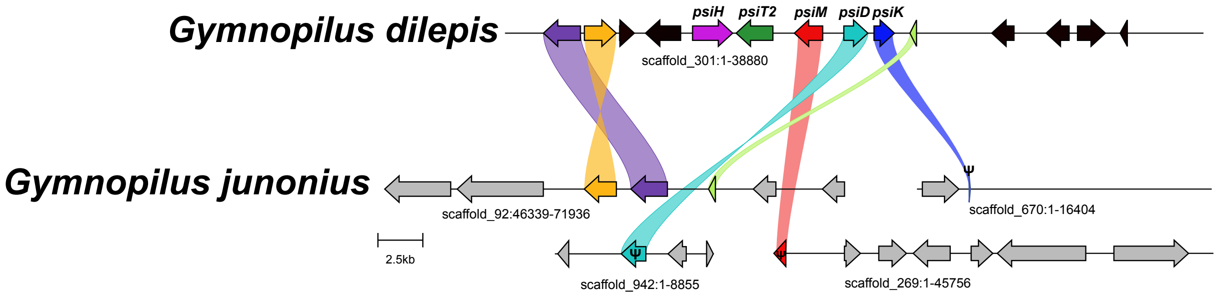


**Figure S2. Evidence of loss of the PGC in *Gymnopilus junonius*.** Three pseudogenes were recovered by tblastn and respective scaffolds were aligned using clinker and clustermap.

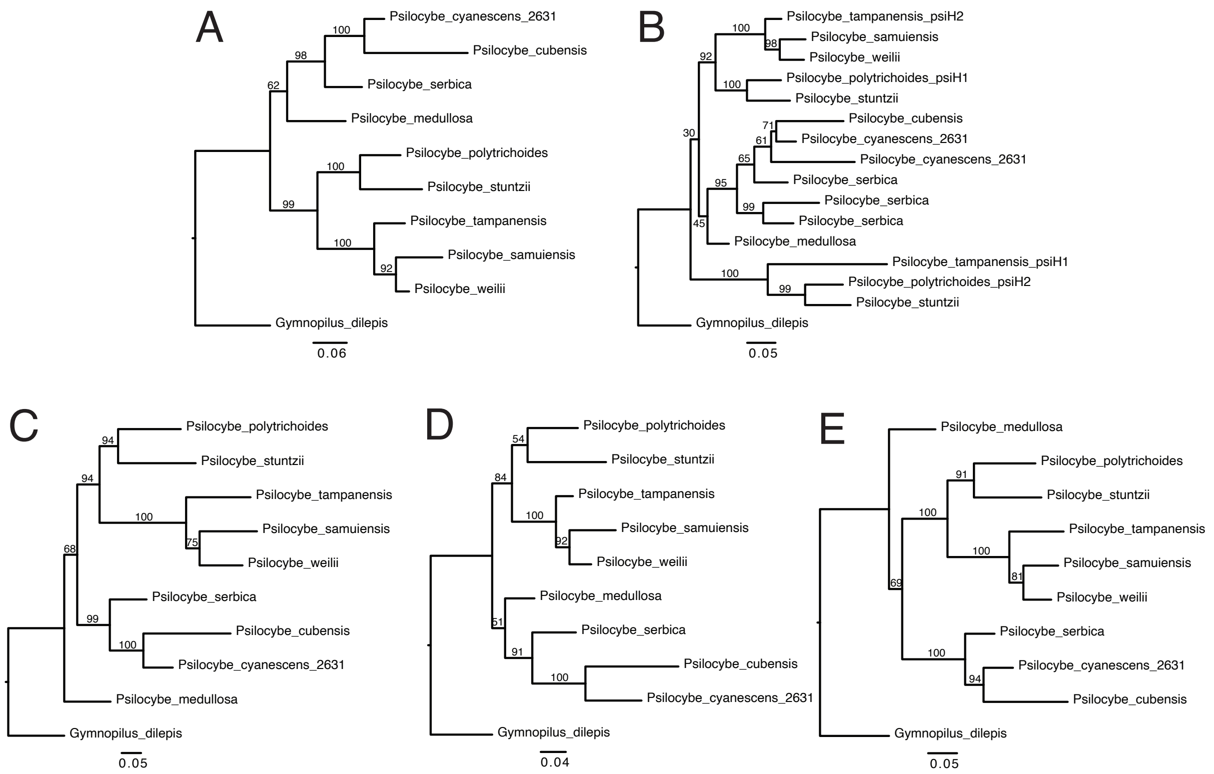


**Figure S3. Phylogenetic analyses of PGC sequences weakly place Clade II within Clade I.** Maximum likelihood phylogenies of PGC amino acid sequences. A. PsiD, B. PsiH, C. PsiK, D. PsiM, E. PsiT2, with support values representing percentage of 1000 ultrafast bootstraps. All gene topologies are consistent with HGT from Clade I to Clade II of *Psilocybe*, but this assumes *Gy. dilepis* represents a true outgroup.

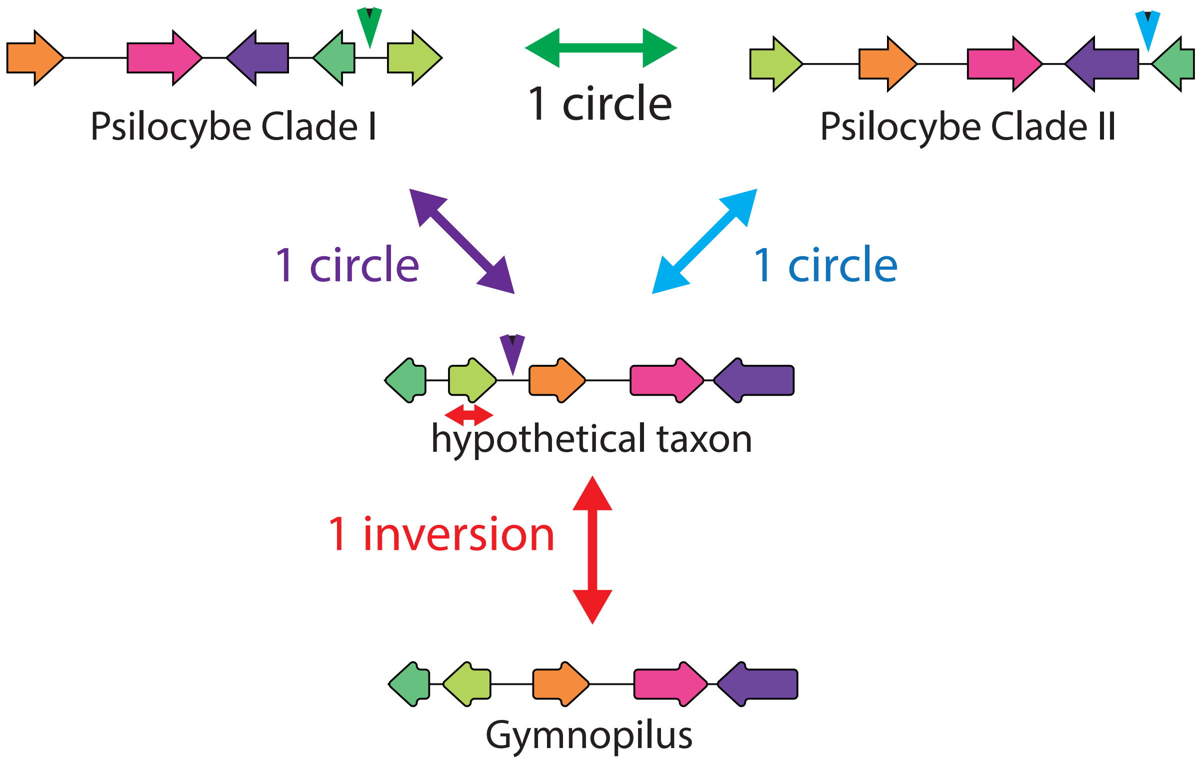


**Figure S4. Molecular evolution events explaining the diversity of PGC gene order.** One circular intermediate separates the gene order of *Psilocybe* Clade I, Clade II, and a hypothetical intermediate taxon, which differs from the PGC order in *Gymnopilus* by a single gene inversion. Breakpoints are indicated by arrowheads whose color corresponds to the circular intermediate, and the inversion is indicated by a bidirectional arrow.
